# Cell-type specific patterned stimulus-independent neuronal activity in the *Drosophila* visual system during synapse formation

**DOI:** 10.1101/376871

**Authors:** Orkun Akin, Bryce T. Bajar, Mehmet F. Keles, Mark A. Frye, S. Lawrence Zipursky

## Abstract

Stereotyped synaptic connections define the neural circuits of the brain. In vertebrates, stimulus-independent activity contributes to neural circuit formation. It is unknown whether this type of activity is a general feature of nervous system development. Here, we report patterned, stimulus-independent neural activity in the *Drosophila* visual system during synaptogenesis. Using *in vivo* calcium, voltage, and glutamate imaging, we found that all neurons participate in this spontaneous activity, which is characterized by brain-wide periodic active and silent phases. Glia are active in a complementary pattern. Each of the 15 examined of the over 100 specific neuron types in the fly visual system exhibited a unique activity signature. The activity of neurons that are synaptic partners in the adult was highly correlated during development. We propose that this cell type-specific activity coordinates the development of the functional circuitry of the adult brain.

## Introduction

Synaptic connections between neurons determine how neural circuits process information. Understanding how the specificity of these connections is established is a central challenge in neurobiology. In vertebrates, cell autonomous genetic programs and neural activity—both evoked and spontaneous—contribute to the development of synapses. Spontaneous activity has been observed throughout the developing central nervous system (CNS)—in the hippocampus (Ben-Ari et al., 1989), spinal cord (Landmesser and O’Donovan, 1984), cerebellum (Watt et al., 2009), auditory system (Tritsch et al., 2007), and visual system (Galli and Maffei, 1988; Meister et al., 1991). Retinal waves were discovered over 20 years ago and are the best characterized examples of spontaneous activity (reviewed in Ackman and Crair, 2014; Blankenship and Feller, 2009; Kirkby et al., 2013; Sernagor and Hennig, 2013). In mice and other mammalian models, retinal waves begin soon after the completion of axon guidance and persist through eye opening. During this period, bursts of activity propagate from the retina to higher visual centers, including the lateral geniculate nucleus (LGN), the superior colliculus (SC), and the visual cortex (Ackman et al., 2012). In each of these areas, large populations of neighboring cells exhibit correlated firing patterns. Significant progress has been made toward characterizing and identifying the organizing principles of spontaneous activity in the developing vertebrate brain, and the precise developmental role of this activity is an area of active interest.

By contrast to vertebrates, brain development in invertebrates is thought to be driven by hardwired morphogenetic programs driven by cell recognition molecules, with little role for spontaneous or experience-dependent neural activity. Previous work has shown that, in the *Drosophila* visual system, photoreceptor neurons can develop the wild-type complement of synapses in a stimulus-independent manner (Hiesinger et al., 2006). However, the existence and significance of spontaneous activity during invertebrate brain development remains an open question.

Some of the most detailed understanding of brain development in the fly comes from the visual system. Visual information from the compound eye is relayed in a topographic fashion to the optic neuropils—the lamina, medulla, and the lobula complex. These neuropils are organized into columns and layers. In general, columns process information from different points in visual space, and layers process different types of visual information. Over 100 different neuronal cell types form precise synaptic connections, typically with several different cell types. The three dimensional EM re-constructions of the optic neuropils that reveal this wiring complexity (Rivera-Alba et al., 2011; Takemura et al., 2013, 2017) also underscore the challenge of understanding the mechanisms of synaptic specificity: Most neurons make synapses with only a subset of their contact neighbors, and the area of contact has little bearing on this decision.

Visual system development in the fly takes place during the last stage of larval development and the ensuing 100 hours of metamorphosis, or pupal development. Synapse formation, as well as axon guidance and morphogenesis, are predicated on cell-cell contacts. As such, much of the focus in the study of neural development has been on the roles of cell surface and recognition molecules. This body of work, carried out at the level of individual cell types, paints the picture of a dynamic self-assembly process in which local interactions shape the developmental trajectory of each neuron (Hadjieconomou et al., 2011; Huang et al., 1998; Pecot et al., 2014). By 50 hours after pupa formation (hAPF), these specific and genetically hardwired molecular push-pulls bring most of the cell types of the visual system to where they belong in the adult brain, ready for synaptogenesis. Over the remaining 50 hours of pupal development, synapse assembly proceeds in parallel with notable changes in gene expression, including the upregulation of genes involved in neural activity and new sets of cell recognition molecules (Chen et al., 2014; Tan et al., 2015; Zhang et al., 2016). It is during this time that vast networks are assembled, comprising distant cells which must be linked through specific synaptic connections and compatible gene expression profiles (e.g. matching neurotransmitter systems and receptors). Little is known about the molecules and mechanisms that coordinate this period of brain development.

Here we report the discovery of stimulus-independent neural activity in the developing *Drosophila* CNS and its initial characterization in the visual system. We find that the visual system as a whole, all 15 of the individual neuronal cell types examined, as well as astrocytic glia, participate in <u>p</u>atterned, <u>s</u>timulus-<u>i</u>ndependent <u>n</u>eural <u>a</u>ctivity (PSINA), during the late stages of circuit assembly. We speculate on the function of this globally organized, cell type-specific activity in regulating the development of the connectome.

## Results

### Patterned neuronal and glial activity in the developing fly brain

To assess whether neural activity contributes to visual system development in *Drosophila*, we used an *in vivo* 2-photon live imaging protocol that enables continuous observation over several days (i.e. from ~16 hrs after puparium formation (hAPF) to eclosion) (Akin and Zipursky, 2016; Langen et al., 2015). We expressed the genetically-encoded calcium indicator (GECI) GCaMP6s (Chen et al., 2013) throughout the central nervous system using a pan-neuronal GAL4 driver. Between 40 and 50 hAPF, the optic lobe is largely inactive, aside from sporadic activity in isolated cells or groups of cells with no discernable spatial or temporal coordination (Movie S1). Starting shortly before 50 hAPF, a subset of neuronal processes begins to exhibit periodic pulse trains of increased fluorescence. By 55 hAPF, neuronal processes in all optic neuropils, as well as fibers originating from the central brain, participate in regular 12-15 minute long cycles of active and silent phases (**Figures 1A-1B, Movie S1**).

**Figure 1.**
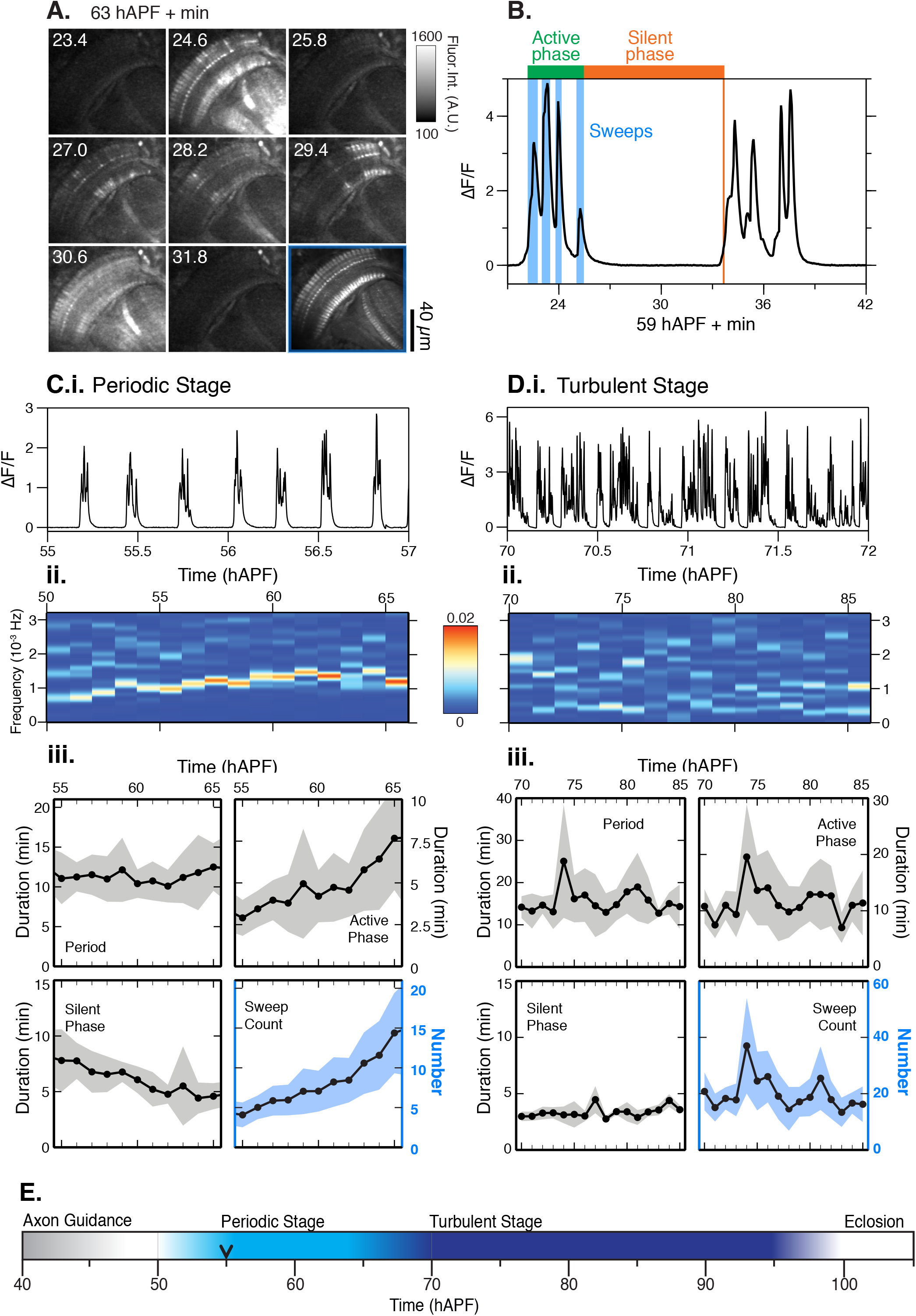
Patterned stimulus independent neural activity (PSINA) in the developing visual system. **A.** Micrograph montage showing a single cycle at 63 hAPF; framed panel (lower right) is the average intensity projection through the active phase. **B.** Representative cycle showing four sweeps (duration indicated in blue) during a shared active phase (green), and a shared silent phase (orange). **C.i.** Representative trace of a 2 hr interval during the periodic stage (50-65 hAPF). **C.ii.** Frequency analysis (Fourier transform) between 50-65 hAPF; **C.iii.** Average traces of cycle metrics in the periodic stage (n = 54 ROIs from 6 flies). Shaded areas represent standard deviation. **D.i.** Representative trace during the turbulent stage (70 hAPF to eclosion); **D.ii.** Frequency analysis (Fourier transform) between 70-85 hAPF; **D.iii.** Average traces of cycle metrics in the turbulent stage (n = 46 ROIs from 4 flies), shaded areas represent standard deviation; **E.** Summary of spontaneous activity stages during pupal development. Black arrowhead marks the time point after which 100% of columns participate in each cycle. See Table S1 for genotypes used in this figure.

The active phase of each cycle comprises several distinct bouts of activity, each of which may have one or a set of closely spaced peaks (**Figure 1B**). We term these bouts *sweeps.* Between 55 and 65 hAPF, the cycle period remains roughly constant while the number of sweeps per cycle and the duration of the active phase increase at the expense of a shrinking silent phase (**Figure 1C**). Fourier analysis captures the simple periodicity of the activity pattern with a single dominant band for the cycle frequency (~0.001-0.002 Hz) over what we call the *periodic stage* (**Figure 1C**). There were no significant differences between periodicity, sweeps per cycle, active and silent phase durations between different animals, consistent with the notion that the developmental mechanisms underlying these metrics are stereotyped (**Figure S1A**).

To further characterize the time evolution of the neuronal activity, we moved our expression system to the *cn,bw* genetic background (Thimann and Beadle, 1937), which eliminated pigmentation in the retina and allowed us to image through eclosion. We found no significant differences in the periodic stage activity metrics between *cn,bw* flies and control flies (**Figure S1A**). Imaging beyond this stage revealed that by 70 hAPF the earlier, simple temporal pattern is replaced with multiple frequencies reflecting cycles with variable periods (**Figure 1D, Movie S1**). Compared to the periodic stage, during this later, *turbulent stage,* individual cycles exhibit higher sweeps per cycle, and, on average, longer and shorter active and silent phase durations, respectively (**Figure 1D**). The turbulent stage persists until the final hour of pupal development, after which the number and amplitude of peaks drop before eclosion (**Figure S1B**). Thus, activity during development is divided into an early periodic stage and a later turbulent stage, and continues until an hour before the adult fly emerges (**Figure 1E**).

We next asked whether activity was present beyond the visual system. Recently, a detailed study of motoneuron development showed that the neurons of the peripheral nervous system exhibit periodic bouts of activity, starting at 48 hAPF, which grow stronger as development proceeds (Constance et al., 2018). In pupae, the pan-neuronally driven GCaMP6s was sufficiently bright to detect using a wide field-of-view epifluorescence microscope, making it possible to image the whole CNS of multiple animals simultaneously. We used this alternative preparation to follow the GECI signal between 58 and 60 hAPF, and observed cycles of activity that matched our observations from the 2-photon setup in both the optic lobes and the central brain (**Figure 2A, Movie S2**). We also established that the pattern of activity is the same in males and females (**Figure S2A**).

**Figure 2.**
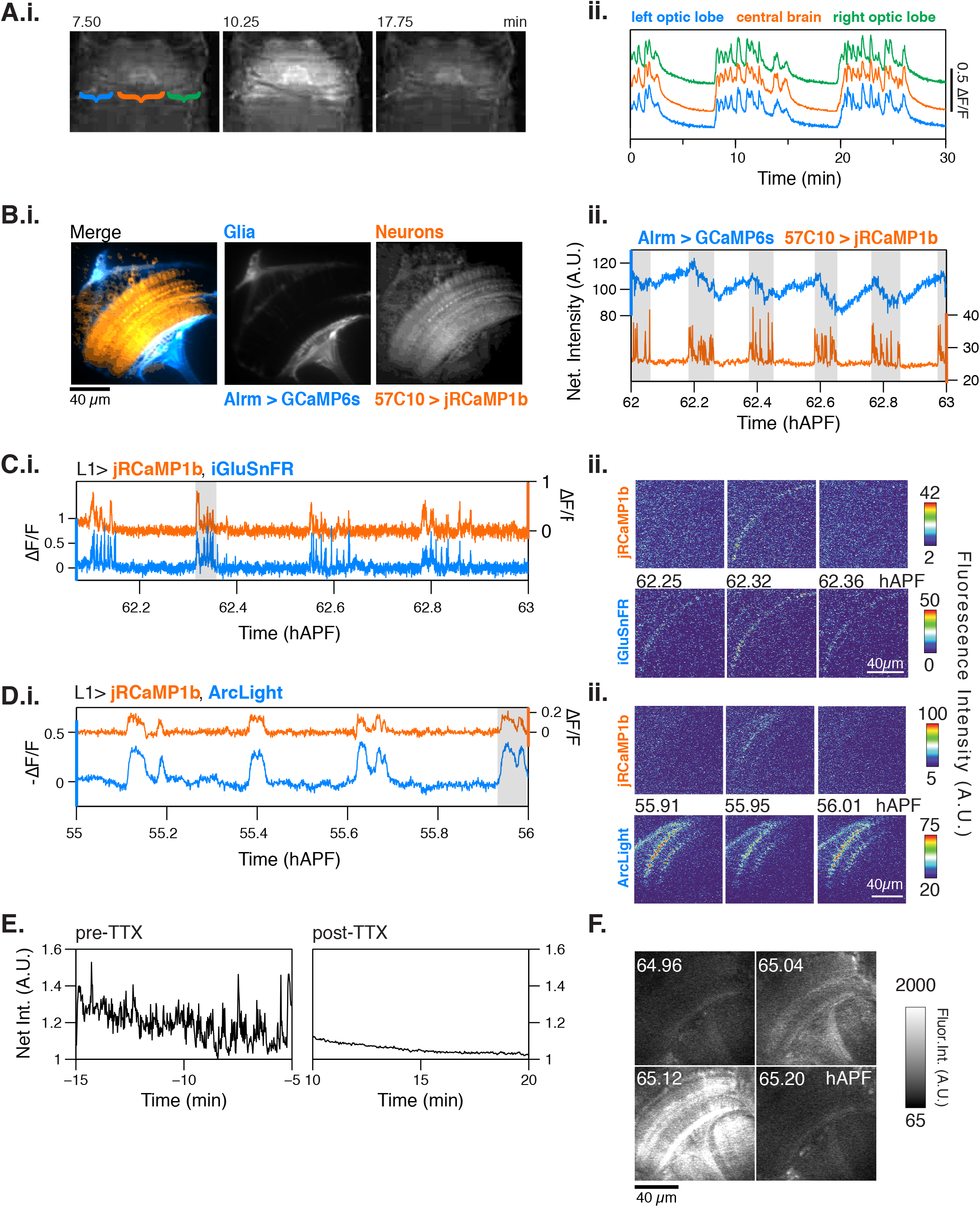
Characterization of PSINA. **A.i.** Representative epifluorescence images of a single cycle in an intact pupa expressing pan-neuronal GCaMP6s. Brackets mark left optic lobe (blue), central brain (orange), and right optic lobe (green). **A.ii.** Average traces from ROIs encircling the left optic lobe (blue), central brain (orange), and right optic lobe (green) between 58-60 hAPF. **B.i.** Representative micrograph showing astrocytic glia expressing GCaMP6s (blue) and pan-neuronal expression of jRCaMP1b (orange). Scale bar, 40 μm. **B.ii.** Representative trace comparing glial (blue) and neuronal activity from (orange) between 62-63 hAPF. Active phases of the neuronal cycles are shaded in gray. **C.** Representative traces (**i.**) and micrographs (ii.) from L1 neurons expressing jRCaMP1b (orange, top) and iGluSnFr (blue, bottom). Note that iGluSnFr reports more sweeps than jRCaMP1b; we suspect that this is due to the L1-expressed glutamate sensor’s response to neurotransmitter released by L1 itself, neighboring cells or both. **D.** Representative traces (**i.**) and micrographs (**ii.**) from L1 neurons expressing jRCaMP1b (orange, top) and ArcLight (blue, bottom). **E.** Representative traces of activity as reported by pan-neuronal GCaMP6s before (left) and after (right) addition of 1μM TTX. **F.i.** Micrographs of *norpA*^null^ mutant flies expressing pan-neuronal GCaMP6s shows that visual stimuli are not required for activity to occur. See Table S1 for genotypes used in this figure.

Given the broad domain of the activity, we assessed whether the glial complement of the CNS also participate in this process. Using orthogonal expression systems, we expressed GCaMP6s in astrocytic glia and RCaMP1b in all neurons (**Figures 2B, S2B-D**). Prior to 55 hAPF, there is no significant correlation between the glial GECI signal and neuronal activity (**Figure S2B**). This changes markedly further into the periodic stage when the glial signal begins periodic oscillations (**Figures 2B and S2C**). While, by contrast to the rapid neuronal responses, the changes to the glial GECI signal are tonic, these cells also cycle through periods of high and low intensity alongside neurons, albeit with a notable phase shift: when neurons are active, astrocytes exhibit a progressive loss in GECI signal which is rebuilt during the neuronal silent phase (**Figures 2B and S2D**).

As flies leave the field of view of the microscope upon eclosion, it was not possible to establish whether the oscillatory activity ceases altogether in the adult. To address this question, we used a head-fixed cranial window preparation of adult flies expressing GCaMP6s pan-neuronally (Aptekar et al., 2015; Seelig et al., 2010). We observed stimulus-independent activity in newly eclosed (1 hr old) flies as well as 1- and 5-day old adults, all of which also had intact, robust responses to visual stimuli (**Figure S3A**). By contrast to what we observe during development, spontaneous activity in the adult did not engage the entire optic lobe, exhibited fewer sweeps in a given cycle and oscillated at a higher frequency (**Figure S3B**). Further, with increasing age, less activity was observed, suggesting that the mechanisms driving this regime of activity decreased over a period of days. These differences suggest the involvement of different molecules and mechanisms in generating the pupal and adult stimulus-independent activities.

### Calcium activity correlates with changes in membrane voltage and neurotransmitter release and is independent of visual stimulus

The clear separation between stimulus-dependent and spontaneous GECI signals in the adult, and the interpretation of the former as evoked neuronal activity, raises two questions about the pulses observed during metamorphosis. First, are the pulses we observe during development reporting the electrical excitation across neurons, or rather are they indicative of membrane activity-independent modulation of intracellular calcium levels? And second, if the pupal pulses in the visual system do indeed reflect neuronal activity, do they depend on visual stimuli and phototransduction?

To address whether the pupal pulses reflect neuronal activity, we first asked if the GECI signal is accompanied by temporally matched neurotransmitter (e.g. glutamate) release and changes in membrane voltage by co-expressing the red-shifted GECI, RCaMP1b (Dana et al., 2016), with either the genetically encoded glutamate sensor iGluSnFR (Marvin et al., 2013) or the genetically encoded voltage indicator ArcLight (Jin et al., 2012). Pan-neuronal co-expression of both indicator pairings revealed glutamate release and membrane voltage dynamics that were closely correlated with the GECI signal cycles (**Figures S4A-B**). As discussed below, all neuronal cell types we studied individually also display the same GECI reported activity pattern we describe for pan-neuronal expression. As such, we constrained the source of the co-expressed indicators to a single cell type, the L1 lamina monopolar neuron, a glutamatergic first-order interneuron (Gao et al., 2008; Takemura et al., 2011). With L1, we observed strong correlation between the GECI signal and both the iGluSnFR-reported glutamate release (**Figure 2C, Movie S3**) and ArcLight-reported drops in membrane voltage (**Figures 2D, Movie S4**) at the level of individual sweeps.

To further examine the nature of the pupal pulses, we used the head-fixed cranial window preparation in late stage pupae (90-95 hAPF) to enable pharmacological manipulations. We found that on-going calcium activity at this stage is severely attenuated with the administration of tetrodotoxin, a voltage-gated sodium channel blocker that inhibits action potentials (**Figures 2E and S4C**). Together, these results indicate that the GECI signal observed during pupal development reflects neuronal electrical activity.

Next, we assessed the contribution of visual stimulus to developmental neuronal activity by following the GECI signal in *norpA*^nall^ animals. *NorpA* encodes phospholipase C, which is required in photoreceptors to initiate signaling downstream of rhodopsin, the light-activated G-protein coupled receptor (Bloomquist et al., 1988). We confirmed that *norpA*^null^ animals are blind by testing the optomotor response to wide-field stimulus and closed-loop bar fixation (**Figure S5A-C**). Returning to our calcium imaging preparation, we found that activity was still present during pupal development (**Figure 2F**), although the cycle period was altered (**Figure S5D**). The finding that this activity is independent of visual stimuli is consistent with the reported timing for the onset of photoreceptor light response at 82 hAPF, which is some 27 hours after the neuronal activity begins (Hardie et al., 1993). It is also consistent with our experimental setup in which the only light source is the pulsed IR laser used for 2-photon excitation, which is outside of the sensitivity spectrum of fly photoreceptors (Salcedo et al., 1999). We conclude that the pupal neuronal activity is independent of a visual stimulus. In the remainder of the text we refer to this phenomenon as <u>p</u>atterned, <u>s</u>timulus-<u>i</u>ndependent <u>n</u>euronal <u>a</u>ctivity, or PSINA. Taken together, our observations indicate that PSINA is a globally coordinated process that involves the entire developing CNS.

### Cell type-specific dynamics of PSINA

We returned to 2-photon imaging of the developing visual system to assess PSINA in specific neuronal types. Using GCaMP6s, we followed calcium activity in 15 cell types, representing some of the major visual system classes (i.e. photoreceptors (R7, R8), lamina monopolar neurons (L1, L3, L5), medulla intrinsic neurons (Mi1, Mi4), distal medulla neurons (Dm3, Dm4, Dm9), transmedullary neurons (Tm3, Tm4, Tm9), and T neurons (T4, T5)) (**Figure 3A**). Between 50 and 65 hAPF, the temporal pattern of PSINA in all neurons closely followed the pan-neuronal archetype of the periodic stage; as a group, cells of a type cycled through active and silent phases lasting 12-15 minutes, starting around 50-55 hAPF and gradually increasing the duration and the sweep complement of the active phase over time (**Figure 3B**).

**Figure 3.**
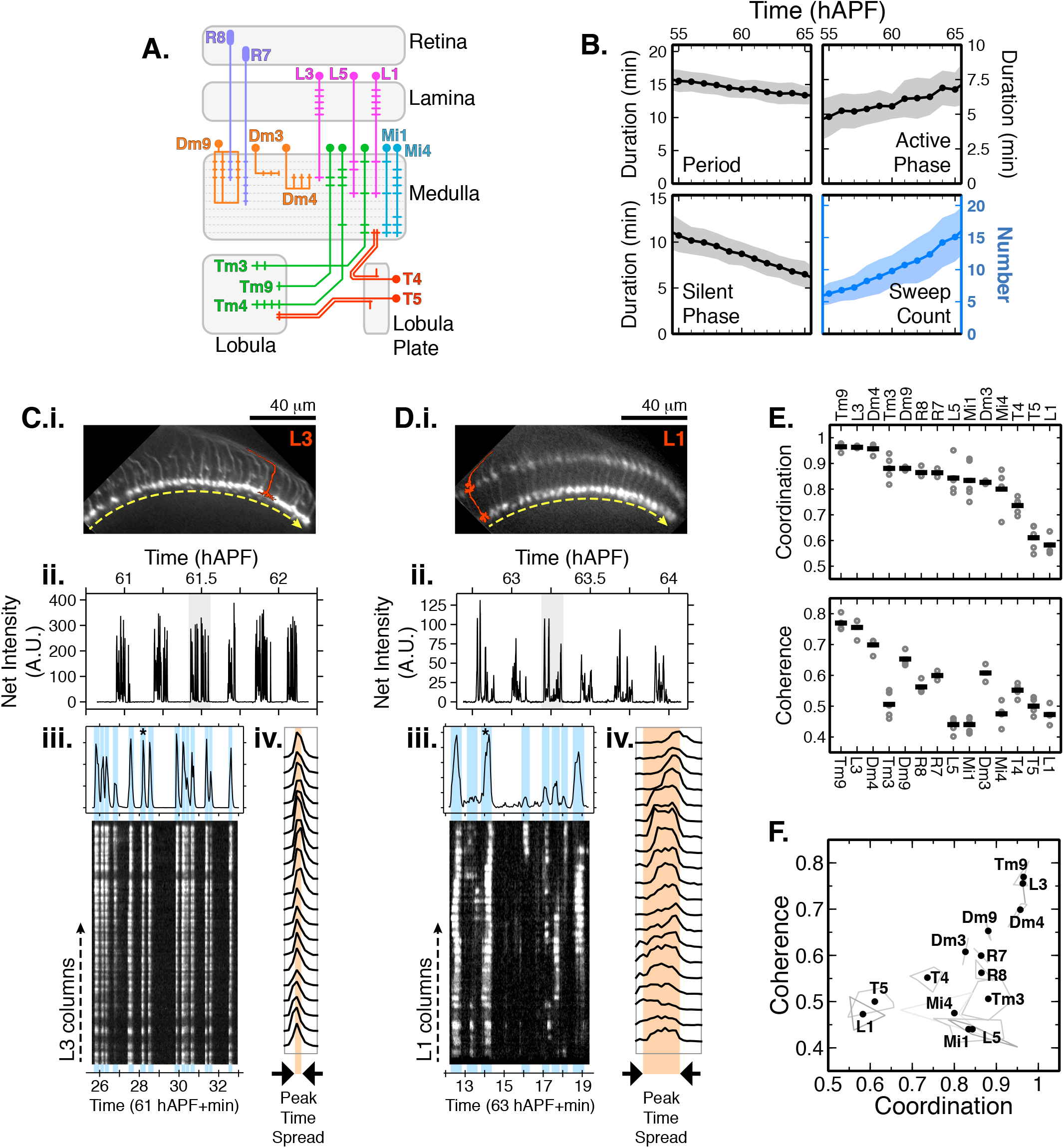
Cell type specific PSINA dynamics. **A.** Schematic of visual system cell types described in Figures 3 and 4; adapted from (Strother et al., 2017) **B.** Cycle metrics in the periodic stage, averaged over 15 cell types and 55 time series. Shaded areas, standard deviation. **C.** PSINA dynamics in L3 cells. **C.i.** Average intensity projection of GCaMP6s expressing L3 processes in the M3 layer of the medulla neuropil. Single L3 schematically shown in red. Dashed yellow arrow sits below the thin profile through M3 used to generate the kymograph in (iii); direction matches the layout of the columns in the kymograph. **C.ii.** Average net fluorescence intensity along the profile described in (i). Active phase with gray background shown in greater detail in (iii). **C.iii.** Plot shows expanded view of an active phase with sweeps highlighted in light blue. Star marks the sweep expanded into individual column traces in (iv). Kymograph of net fluorescence derived from the profile described in (i). **C.iv.** Plot of fluorescence change in individual medulla columns in the star marked sweep in (iii). **D.** Same as (C) for an L1 time series. Kymograph generated from a thin profile through the L1 processes in M5 (i.e. layer just above the yellow line). **E.** Coordination (top) and coherence (bottom) values calculated for different cell types. Round gray markers are individual time series, black bars are the average for each cell type. Between two and six time series shown for each cell type. Metrics for each time series calculated over 55-65 hAPF, using an average of 41+/-9 cycles and 10-20 columns per cycle. **F.** Scatter plot of coordination v. coherence. Vertices of light gray polygons, individual time series; black dots, average for each cell type. See Table S1 for genotypes used in this figure.

**Figure 4.**
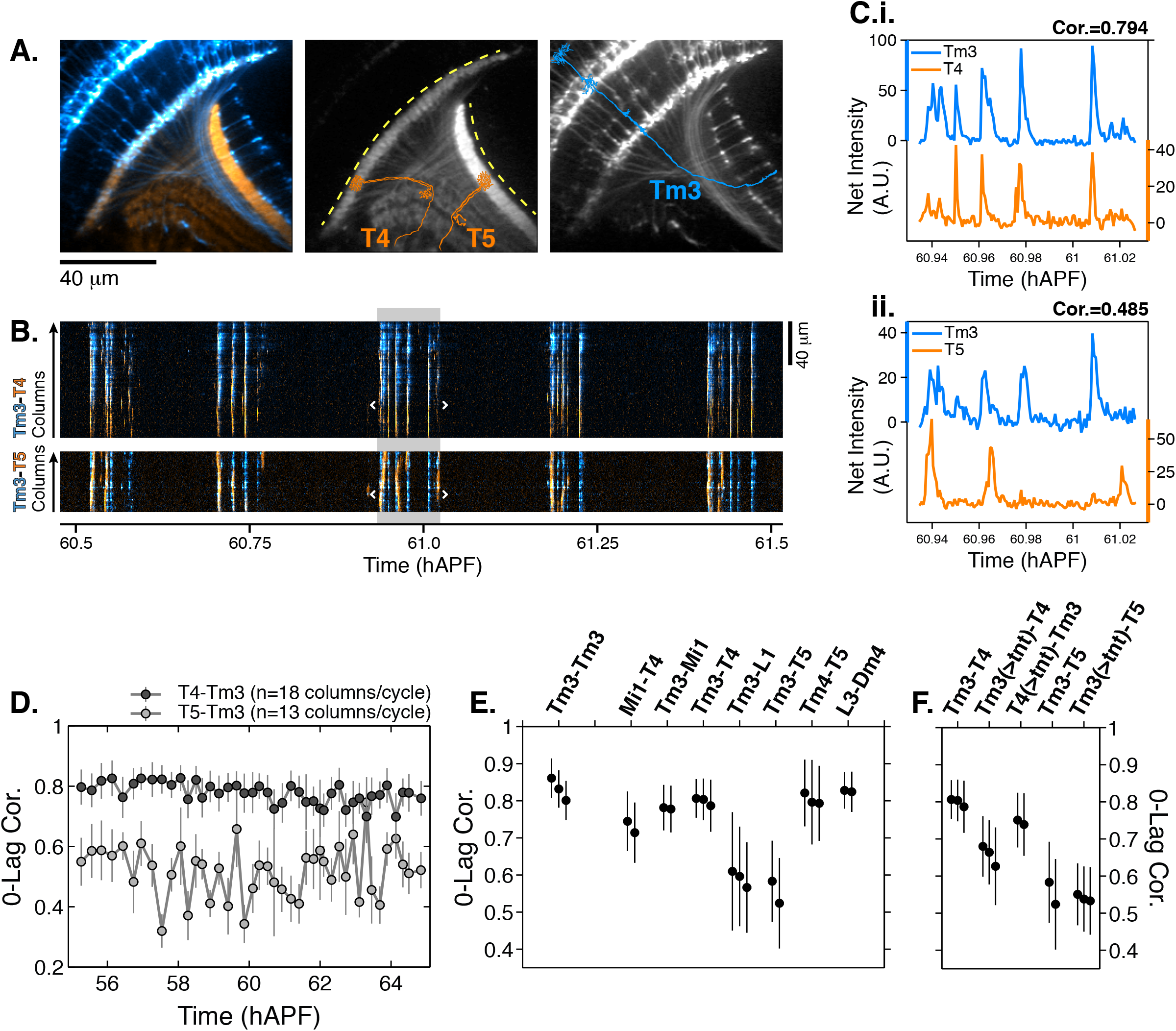
Synaptic release is required for correlated PSINA activity. **A.** Average intensity projection images of GCaMP6s expressing Tm3 (blue) and RCaMP1b expressing T4-5 (orange) cells. Single Tm3, T4, and T5 projections are schematically shown in blue and orange. Dashed yellow arcs in center panel abut the thin profiles through M9-10 and the lobula used to generate the kymographs in (B). **B.** Tm3-T4 (top) and Tm3-T5 (bottom) kymographs of net fluorescence derived from the profiles described in (A). Columns between the white brackets in active phase with gray background were used to generate the plots in (C). **C.** Tm3-T4 (i) and Tm3-T5 (**ii**) net fluorescence intensity along the columns marked in (B). **D.** 0-Lag cross correlation values between 55-65 hAPF for Tm3-T4 (dark gray) and Tm3-T5 (light gray) for the time series used in (A-C). Markers are the average correlation value for 10-20 columns per cycle, gray vertical lines are standard deviation. **E.** 0-lag correlation values for pairs of cell types, averaged over 55-65 hAPF. Black markers and vertical lines are the average and standard deviation for each time series. 2-3 time series shown per pair. 43+/-10 cycles with 15+/-3 columns per cycle used for each time series. The Tm3-Tm3 pair represents the highest correlation we expect to observe for a perfect match given the signal-to-noise statistics of the data (see Figures S6A-S6B). **F.** TNT expression in Tm3 reduces Tm3-T4 correlation but has no effect on Tm3-T5 correlation. Data statistics as in (E). Unperturbed pairs reproduced from (E) for ease of comparison. See Table S1 for genotypes used in this figure.

Whereas the broad temporal characteristics of PSINA are shared between all neuronal types, how the activity propagates across the repeated columnar array of a given cell type varies significantly. For example, nearly all L3 neurons participate in every sweep of an active phase while in L1s fractional participation can change notably between sweeps (**Figures 3C and 3D, Movie S5**). Further, during a sweep, L3s reach peak intensity within narrower time window compared to L1s (**Figures 3C and 3D**). In order to compare PSINA dynamics between repeated observations of the same cell type and across different cell types, we defined two scalar metrics, *coordination* and *coherence,* to represent the distributions of fractional participation and peak time spread values, respectively. Coordination is the fraction of sweeps with greater than 90% column participation. Coherence is the largest fraction of columns that peak within the same time point, averaged over all sweeps. Accordingly, distinct observations of PSINA in L3s all yield comparably high coordination and coherence values in contrast to L1, which scores consistently lower for both metrics (**Figure 3E**). We extended this analysis to 13 other cell types and found that coordination and coherence values from separate observations cluster around means characteristic to each cell type, independent of the specific drivers used for GECI expression or transgenic constitution of the animals (**Figures 3F and S6A, Movie S6**). For most cell types, coordination and coherence are roughly constant during the periodic stage, between 55-65 hAPF (**Figure S6B**). In the few that do show changes, we observe loss of coordination that is attributable to loss of image quality as developing retinal pigmentation degrades the observed GECI signal, particularly with weaker cell type-specific drivers. A notable exception is L1; here, despite the loss of net signal, both metrics increase over time (**Figure S6B**), indicating that the observed trends reflect evolving PSINA dynamics. Visual inspection of the L1 activity pattern over time confirms this conclusion (**Figures S6C-S6D**). In summary, we find that the fine spatio-temporal structure of PSINA is cell type-specific, stereotyped and can be dynamic over the course of development.

The cell type-specific diversity of PSINA dynamics suggested that the activity may be traversing the visual system through multiple parallel ‘channels’ which engage distinct sets of neurons. To explore this notion, we imaged pairs of neurons expressing red and green GECIs (**Figure 4**). For example, we compared the activity in Tm3 transmedullary neurons, with processes in both the medulla and the lobula neuropils, to the medulla-resident dendrites of the T4 class and the lobula-resident dendrites of T5s (**Figures 4A-4D, Movie S7**). Between 55-65 hAPF, the Tm3-T4 activity was highly correlated (0.8+/-0.06, n=3) while the Tm3-T5 correlation was significantly lower (0.55+/-0.1, n=2) (**Figures 4D-4E, S7A-S7B**). The results were the same when these measurements were repeated with the opposite cell type and color pairing (**Figure S7C**). Notably, in the adult, Tm3 and T4 are synaptic partners in the ON-motion circuit (Takemura et al., 2013) while T5, which is part of the OFF-motion circuit, is not a synaptic partner with Tm3 (Shinomiya et al., 2014).

Downstream of photoreceptors, L1 is considered to provide the principal input to the ON-motion circuit (Joesch et al., 2010), with Tm3 and Mi1 as its major post-synaptic partners, which then synapse with T4 (Behnia et al., 2014; Takemura et al., 2013), the first direction selective neuron in the pathway (Maisak et al., 2014) (**Figure 3A**). We found that the activities of the Mi1-Tm3 and Mi1-T4 pairs are also well correlated while L1-Tm3 has lower correlation (**Figure 4E**). As discussed above, the dynamics of PSINA in L1 evolve through pupal development, and may eventually converge with a presumptive ON-motion PSINA channel during the ensuing turbulent stage. Alternatively, if PSINA is propagated through some form of synaptic coupling, the low L1-Tm3 correlation may be reporting on the sign of the interaction; that is, L1 could be an inhibitory synaptic partner at this stage of development.

Returning to T5, we found that its activity is highly correlated with Tm4, an OFF-motion circuit input into T5 in the adult (Serbe et al., 2016; Shinomiya et al., 2014) (**Figure 4E**). Finally, we observed highly correlated activity between a pair of high coordination-coherence cells, L3 and Dm4 (**Figure 4E**), which are also synaptic partners in the adult. Together, these results confirm the presence of multiple distinct channels of PSINA activity.

Correlated activity patterns between many adult synaptic partners some 45 hours before the end of pupal development hinted at the existence of an early form of synaptic pairing. To explore whether the observed correlations depend on synaptic release, we expressed tetanus toxin (TNT) in one cell type of a pair and measured the correlation of the PSINA activities as before. Driving TNT expression in Tm3s reduced the correlation of the Tm3-T4 pair significantly while the Tm3-T5 value was unaffected (**Figure 4F**). By contrast, T4 expression of TNT had no effect on the Tm3-T4 correlation (**Figure 4F**). These results indicate that the coordinated PSINA activity in Tm3 and T4 is dependent on synaptic release from Tm3, consistent with the notion that PSINA propagation is achieved through some form of synaptic coupling.

## Discussion

In summary, here we report the discovery and initial characterization of PSINA in the developing fly visual system. We observe three distinct stages of PSINA: a periodic stage between 55 and 65 hAPF, a turbulent stage lasting from 70 hAPF to the final hour of pupal development, and an adult stage that persists alongside mature stimulus responses through at least the first four days following eclosion. During the periodic stage, which coincides with the onset of synaptogenesis in the fly CNS (Chen et al., 2014; Muthukumar et al., 2014), each neuronal cell type of the 15 analyzed exhibited stereotyped and distinct activity patterns. Many adult synaptic partners had correlated activity, which depended on synaptic release.

Distinct periodic calcium dynamics were also seen in astrocytes. Astrocytes in the developing adult brain elaborate processes which infiltrate the neuropil during synapse formation (Muthukumar et al., 2014). Ablating astrocytes leads to a significant reduction in the total synapse count (32-47%, depending on the region) in the brain, supporting a role for these cells in regulating synaptogenesis (Muthukumar et al., 2014). Astrocytes of the optic neuropils also elaborate their processes over the same time period (Richier et al., 2017). Here we report that astrocytes exhibit cycles of GECI signal that are matched, though offset, to the periodic PSINA. These findings raise the possibility that astrocytes, spontaneous activity in them, and PSINA contribute to the formation, specificity, or maturation of synapses within the visual system.

What is the contribution of PSINA to building a brain? The best characterized system are retinal waves, which drive the activity of retinal ganglion cells (RGCs), the exclusive conduit of information from the eyes to the brain. Here, RGC projections from both eyes target the LGN and the SC, where they create retinotopic maps of the visual field and segregate based on the eye of origin. In the mouse, retinal waves are described in three stages: The gap-junction mediated stage I from embryonic day 17 (E17) to post-natal day 1 (P1), the ‘cholinergic’ stage II between P1-P10, and the ‘glutamatergic’ stage III from P10 to eye opening at P14 (Blankenship and Feller, 2009; Sernagor and Hennig, 2013). Retinotopy and eye-specific segregation in the LGN and SC are refined over the same period as the second stage of retinal waves. This cholinergic stage is driven by starburst amacrine cells (SACs) (Zheng et al., 2006, 2004). Early work in the field established that pharmacological manipulation of spontaneous activity in the retina disrupts the organization of RGC projections in the LGN (Shatz and Stryker, 1988; Sretavan et al., 1988). Later studies, using progressively more refined methods, have shown that disrupting the cholinergic circuit of SACs and RGCs largely eliminates retinal waves and leads to defects in the refinement of retinotopy and eye-specific segregation of RGC projections (Bansal et al., 2000; Burbridge et al., 2014; McLaughlin et al., 2003; and others). In brief, retinal waves are necessary for the correct patterning of RGC projections in the brain.

Five classes of neurons comprise the retina: Photoreceptors, bipolar cells, amacrine cells, horizontal cells, and retinal ganglion cells (RGCs). The diversity of cell types within these classes—as many as 30 for RGCs (Sanes and Masland, 2015)—is comparable to the fly visual system. Whether there is cell type-specific texture to the retinal waves similar to PSINA described here is not known, although broad classes of RGCs and cone bipolar cells have been shown to exhibit temporally offset firing patterns (Akrouh and Kerschensteiner, 2013; Kerschensteiner and Wong, 2008) With improving genetic handles for distinct retinal cell types and ongoing efforts at describing the high resolution connectome, it will be possible to explore cell type-specific patterns and the contribution of retinal waves to retinal circuitry (Seung and Sümbül, 2014).

In *Drosophila*, peristaltic contractions of body wall muscles have recently been appreciated as part of broad neuronal activity during embryonic development (Vonhoff and Keshishian, 2016). This activity is similar to PSINA observed during pupal development with respect to periodicity and timing relative to synapse formation and refinement. Preventing motor neurons from participating in this neuronal activity, or disrupting calcium-dependent intracellular signaling results in ectopic synapses (Carrillo et al., 2010; Jarecki and Keshishian, 1995; Vonhoff and Keshishian, 2016). In the wildtype, the calcium transients in motor neurons are thought to enable synaptic pruning in response to the muscle-derived chemorepellent Sema2a (Vonhoff and Keshishian, 2016). A similar link between spontaneous activity and axon guidance has also been demonstrated in the developing mammalian visual system. Here, oscillatory Ca^2+^ activity in RGCs were shown to be required for the ephrin-A5 dependent re-positioning of RGC projections in the SC in *ex vivo* cultures (Nicol et al., 2007). These observations from the fly and the mouse suggest that axon guidance and, more broadly, neuronal morphogenesis may be common effectors of spontaneous activity during brain development.

Based on the studies we reference here, previous studies in the fly visual system, and of the role of spontaneous activity in other systems, we propose a general conceptual framework for the role of PSINA in regulating the assembly of the adult connectome. Here, we provide evidence to suggest that some adult synaptic pairings are already established by 55 hAPF, soon after the first pre-synapses can be detected and co-incident with the onset of periodic PSINA. The global coordination of PSINA indicates that an early connectome, one that must be built without activity, is present at this time. This early connectome, comprising the processes of over 100 different neuronal cell types, would be built through local, largely contact-dependent biochemical interactions. While the level of organization achieved through such mechanisms is astonishing, the early connectome may still be a rough approximation of what is required in the adult. PSINA, by orchestrating cellular communication at temporal and spatial scales inaccessible to other signaling mechanisms, may be acting to refine this first draft to complete the self-assembly of the brain. Sweeps of activity repeatedly coursing through the brain through different ‘channels’ could link distinct sets of neurons to direct coordinated morphological changes and sculpt cell-cell contacts, strengthen synapses with correct targets while weakening and pruning incorrect pairings, and control transcription programs that direct circuit refinement (Lee et al., 2017; Nakashima et al., 2013; Serizawa et al., 2006; Tyssowski et al., 2018). PSINA may act as a ‘dress rehearsal’ for neural networks, preparing for ‘opening night’ at the completion of development. Individual cells know their own lines, with whom they interact, and their respective positions on the stage; however, repeated practice of each scene is necessary to refine interactions and ensure that each of the cast can perform as part of a whole ensemble.

We find it remarkable that as a process that appears to engage most, if not all, of the CNS, PSINA is the collective output of the genetically hardwired developmental programs of individual neurons. Thus, despite its complexity, the organizing principles, the driving forces, and the functional significance of PSINA at the level of circuits, cells, and molecules should be discoverable through genetic analysis. Undertaking this effort in the fly visual system, where structures analogous to the vertebrate retinal plexiforms, the LGN, and the SC (Sanes and Zipursky, 2010) are compactly organized in a single microscopic field of view and for which the EM-derived connectome is available, may yield valuable insights into whether and how PSINA affects synaptic specificity and circuit maturation. We expect that the ever-expanding genetic toolkit of *Drosophila,* complemented with improvements in genomic/transcriptomic analysis and imaging technology, will offer a robust experimental track toward understanding PSINA’s contribution to brain development and function.

## Acknowledgments

We thank Gerald M. Rubin, Barrett Pfeiffer, David J. Anderson, Yon-il Jung, Douglas Kim, Vivek Jaramayan, and Yi Sun for GCaMP6s and jRCaMP1b flies, and for technical advice. For assistance and advice with preliminary experiments, we thank Na Ji, Anderson Chen, and Rongwen Lu. For discussing unpublished results and providing feedback on our findings, we thank Kristin Scott, Chris Doe, Marla Feller, and Haig Keshishian. We thank members of the Zipursky Lab for their support and insights. This work was supported by NIH T32 GM008032 (BTB) and NIH R01-EY026031 (MAF), and Howard Hughes Medical Institute. SLZ is an investigator of the Howard Hughes Medical Institute.

## Contributions

BTB and SLZ conceived the original approach. OA, BTB, and SLZ developed the project. OA designed imaging and analysis protocols with assistance from BTB. OA and BTB collected and analyzed developmental imaging data. MFK and BTB collected and analyzed adult imaging and behavior data. OA, BTB, and SLZ wrote the paper. MAF and SLZ supervised the project.

## Supplemental Information

Supplemental Information includes seven figures, one table, and seven movies.

## STAR METHODS

### Key Resources Table

#### Contact for Reagent and Resource Sharing

Further information and requests for resources and reagents should be directed to and will be fulfilled by Orkun Akin.

#### Experimental Model and Subject Details

Flies were reared at 18°C or 25°C on standard cornmeal/molasses medium. Pupal development was staged with respect to white pre-pupa formation (0 hAPF) or head eversion (12 hAPF). The GAL4/UAS and LexA/LexAop expression systems (Brand and Perrimon, 1993; Lai and Lee, 2006) were used to drive cell type-specific transgene expression; complete genotypes used in each experiment can be found in Table S1.

### Method Details

#### 2P Imaging of the Developing Visual System

Pupae were prepared for imaging as previously described (Akin and Zipursky, 2016). Briefly, the cuticle around the heads were removed with fine forceps and the animals were attached eye-down on a coverslip coated with a thin layer of embryo glue. A water reservoir on the objective side of the coverglass provided sufficient immersion medium to last through the hours-long imaging sessions; another reservoir below the pupae kept the animals from dehydrating.

Time-lapse imaging of the visual system was carried out on a custom-built 2P microscope (Akin and Zipursky, 2016) equipped with a 20x water immersion objective (Zeiss, W Plan-Apochromat 10x/1.0 DIC) and 2 GaAsP detectors (Hamamatsu). Over the 2-24 hr imaging sessions, the pupae were kept at 25°C using an objective heater system (Bioptechs). A tunable Ti: Sapphire pulsed laser (Chameleon Ultra II, Coherent) was used as the light source. Green fluors were excited at 940 or 970 nm with ~30 mW under-the-objective power; 1020 nm at ~60 mW was used for red fluors and two-color imaging. Animals imaged under these conditions developed normally and eclosed on schedule. To observe a thicker cross-section of the visual system than possible with a single optical slice, we used the maximum intensity projection of three successive images taken 2 μm apart in the z-axis as the frame for an individual time point. Thus, the effective sampling rate of these time series was 0.4 Hz (2.5 s per frame).

#### Wide-field Imaging

Pupae were staged for head eversion and reared at 25°C. At 58-60 hAPF, pupae were affixed to a Sylgard 184 Silicone Elastomer plate (Dow Corning) with double-stick adhesive tape (3M). Images were acquired with a SteREO Discovery.V8 stereomicroscope (Zeiss) with illumination provided by an X-Cite Series 120PC light source (Excelitas) and captured on a Vixia HF R20 1/4.85 inch CMOS camera (Canon). Images were acquired at 30 Hz. Time series were processed with Fiji (ImageJ) (Schindelin et al., 2012) and analyzed using MATLAB (Mathworks, Natick, MA, USA).

#### Adult Functional Imaging

Calcium imaging was performed as previously described (Keleş and Frye, 2017). Briefly, a single fly was anesthetized at 4°C and placed into a chemically etched metal shim attached to a custom 3D-printed holder. Holder design was based on (Weir et al., 2016); details can be found at http://ptweir.github.io/flyHolder/. The head capsule and thorax were glued to the metal shim using a UV-curable glue (www.esslinger.com). Legs and the antennae were immobilized using beeswax applied with a heated metal probe (Waxelectric-1, Renfert). The head capsule was bathed in saline (103mM NaCl, 3mM KCl, 1.5mM CaCl2, 4mM MgCl2, 26mM NaHCO3, 1mM NaH2PO4, 10mM trehalose, 10mM glucose, 5mM TES, 2mM sucrose) and a small window was opened using fine forceps (Dumont, #5SF). Muscles and fat covering the optic lobe were cleared before placing the fly under a 2P microscope (3i, Denver, CO). Neurons expressing GCaMP6s were imaged at 920 nm using a Ti:Sapphire pulse laser (Chameleon Vision, Coherent). Images were acquired at 10 Hz.

An arena of 48 8×8 LED matrices (470 nm, Adafruit) was used to deliver the visual stimulus. Three layers of blue filter (Rosco no. 59 Indigo) were placed between the screen and the fly to eliminate leakage of the LED light into the PMTs. The screen extended ±108° along the azimuth and ± 72° in elevation. Each LED pixel corresponded to a coverage of 2.2° on the retina equator. However, the projection of each pixel on the retina was variable due to the difference between the curvature of the eye and that of the screen. Visual stimulus consisted of a wide-field grating with a spatial frequency of 35° and presented at a temporal frequency of 0.62 Hz in both directions (ipsi-to-contra and contra-to-ipsi) along the horizontal axis. The presentation order of the visual stimuli was randomized to prevent sensory adaptation. Each experimental condition was tested three to four times per animal.

#### Tetrodotoxin Treatment

Pupal development was staged for white pre-pupa formation and reared at 25°C. Between 90-95 hAPF, the pupal case was removed with fine forceps. These late pupae were prepared for imaging following the protocol described above for adult functional imaging. Viability was verified by leg or trachea movement. Neurons expressing GCaMP6s were imaged at 920 nm using a Ti:Sapphire pulsed laser (Chameleon Vision, Coherent). Images were acquired at 10 Hz.

Tetrodotoxin at 1μM final concentration was mixed into the saline solution after 40 minutes of imaging and the fly was observed for another 20 minutes after the application of the drug. Viability was confirmed before and after tetrodotoxin administration, and the data were excluded from analysis if the animal did not survive the experiment.

#### Visual Flight Simulator

Flies were cold anesthetized at 4°C, tethered to tungsten pins using UV activated glue, and allowed to recover for 1-2 hours in a small, humidified acrylic container with a red desk lamp providing heat. This recovery regime improves flight performance consistency. The majority of the experiments were performed in the afternoon when flies are most active.

A visual flight simulator composed of 32×96 cylindrical green (570 nm) LEDs was used to deliver visual stimuli (Reiser and Dickinson, 2008). The arena covered ± 180° along the azimuth and ± 60° in elevation. Single flies were positioned in the center of the arena and illuminated from above with an 880 nm infrared LED. The shadow cast by the wings was detected with an optical sensor. Signal form this sensor was analyzed by an instrument called the wingbeat analyzer (JFI Electronics Laboratory, University of. Chicago, Chicago, IL, USA) that calculates left and right wing beat amplitudes (WBA). The difference in the left and right WBA is proportional to the fly’s steering effort in the yaw axis.

For bar fixation closed-loop experiments, a dark bar that is 120° in height and 30° in width was presented to the flies. Positional displacement of the bar in the yaw axis was coupled to the steering effort of the fly, allowing the animal to have the active control of the bar position. Each fly was tested for closed-loop fixation behavior for four minutes. To test open-loop optomotor responses, flies were presented with wide-field gratings with a spatial frequency of 30° and a temporal frequency of 3 Hz for four seconds.

#### Quantification and Statistical

##### Analysis Analysis of Pupal Imaging Data

###### Pre-processing

Processing and analysis of image data were carried out with custom scripts written in MATLAB (Mathworks, Natick, MA, USA). Fiji (ImageJ) was used for some user-assisted tasks and figure preparation. Time series were processed in blocks corresponding to ~6 hours of observation (~9000 frames). In the pre-processing step of reducing lateral motion, the general approach of maximizing the cross correlation of individual frames to a reference image was modified to meet the specific challenges of developmental imaging. First, a series of reference images were generated as averages of pools of high signal frames distributed across each block. After ~55 hAPF, the optic lobes begin to twitch with a period similar to that of PSINA. These fast movements can introduce significant blur into the pool-averaged reference images. To reduce this blur, 300 random subsets of each pool were tested to find the sharpest average reference image. Sequential registration of this series of reference images to each other produced a stabilized representation of the visual system which continues to move and grow over the course of observation (Akin and Zipursky, 2016; Langen et al., 2015). In a second step, the registration of the reference series was refined to minimize the movement of a user defined region of interest (ROI). These internally registered reference images then served as local registration targets for nearby frames of the full block. Finally, the block was corrected for any rotational motion of the ROI.

###### Signal and Feature Extraction

Per frame pixel averages of masked regions were used to define raw signal (F) traces from the image time series. Time-dependent fluorescence baseline (F_o_) was estimated using a moving window approach and used to calculate the net signal (F-F_o_, Figures 3 and 4) and change-in-signal ((F-F_o_)/ F_o_, Figures 1 and 2) traces. User-defined, static masks were used for pan-neuronal and glial expression experiments. For cell type-specific experiments, *dynamic masks,* corresponding to the active columns in each cycle, were defined automatically from the kymograph representation of the time series. Briefly, kymographs were generated as concatenated line profiles from user-defined, segmented arcs of 7-9 pixel (3-4 μm) thickness, drawn across a single layer of the medulla or lobula neuropil. Baseline subtracted, net signal kymographs were used in all subsequent analysis. Projecting along the spatial dimension of the kymographs yielded one dimensional net signal traces, which were used to identify the limits of PSINA cycles. Within each active phase, sweeps were defined by ordering intensity peaks with respect to their amplitudes, and, from the largest peak on down, marking the continuous time spans with net signal intensity greater than 75% of peak value; lesser peaks present in the sweep of a larger one were removed from the ordered peak list used in sweep identification. Dynamic masks were based on peaks in the *activity profiles* of PSINA cycles, produced for each active phase by projecting along the temporal dimension of the kymographs. The width of each mask was determined by testing the spatial neighborhood of each peak for correlated net intensity changes in the time domain. The maximum number of dynamic masks identified in each cycle was set to 20.

###### Frequency Analysis

Analysis was implemented in MATLAB, following the guidelines of Uhlen (Uhlén, 2004). Change-in-signal (ΔF/F) traces were processed using a 2-hour sliding window which traversed the time series in 1-hour steps. After filtering with a Hanning window to reduce spectral leakage, each 2-hour block was transformed with the FFT algorithm to obtain non-parametric power spectrum density estimates. The fidelity of the power spectrum density estimate was confirmed by applying the inverse transform on the highest-power peak and comparing the resultant signal to the raw data. One-sided power spectrum density estimates were plotted for each 2-hour block in Figure 1 and Figure S2.

###### PSINA Dynamics

For each cycle, unit signal-to-noise (S2N) value was defined as twice the standard deviation of the net signal trace in the silent phase. A dynamically masked column was considered to participate in a given sweep if it had a net intensity peak greater than or equal to 1.0 S2N within the sweep limits. This scoring scheme was the basis of the definition of the coordination metric. For coherence, the largest fraction of columns that reach peak intensity at the same time point within each sweep was calculated. To ensure consistent comparisons across different cell types, only high participation (≥90%) sweeps were considered for the coherence metric.

###### Correlation Analysis

For the analysis of two-color imaging experiments, two separate kymographs were generated using the same segmented arc. Dynamic masks were derived from the average activity profile of these two kymographs to ensure that the masks captured columns active in both channels. Cycle limits were determined using the brighter channel. For each cycle, masks with a maximum S2N value of at least 1.0 in both channels were used to calculate pairwise 0-lag cross-correlation. Cycles with fewer than 10 masks above the signal quality threshold were excluded in the calculation of time series ensemble statistics (i.e. mean and standard deviation.)

##### Analysis of Adult Calcium Imaging Data

Images were pre-processed to correct for lateral motion using the registration algorithm described above. To find active pixels in the lobula, we defined a mask excluding other neuropils (medulla and lobula plate). For every pixel in this mask, the mean value and standard deviation were calculated for the full time series; the test value for each pixel was defined as the product of these metrics. Pixels with test values greater than or equal to twice the mean value of all pixels in the mask were used in analysis. In our experience, this thresholding approach enriches for active pixels over background and shot noise in the selected mask. The frame average of active pixels were used to produce the signal trace for the time series. Repeated observations were averaged for each fly and a single average trace per experiment was generated.

##### Analysis of Visual Fixation Behavior

Behavioral data from the visual display and the wing beat analyzer was collected with a Digidata 1440A digitizer (Molecular Devices, San Jose, CA, USA) sampled at 1 kHz. Data were processed using custom written scripts in MATLAB (Mathworks, Natick, MA, USA). Briefly, the first 100 milliseconds of the trials were removed and the first data point of the remaining signal was subtracted from the entire trial to set the initial WBA to zero. Delta WBA was calculated by subtracting left from right WBA. Flies which stopped flying during the experiments were excluded from further analysis. Trials for the same experimental conditions were averaged and calculated for all animals. No statistical tests were conducted to pre-determine the sample size. To analyze closed-loop fixation data, the bar position was binned into 96 positions around the visual azimuth and bar histograms for each fly was calculated. Data were then averaged across the animals for the time bar spent at each position.

##### Data and Software Availability

Scripts developed by the authors and used in this study are available upon request.

